# Automated Protein Function Description for Novel Class Discovery

**DOI:** 10.1101/2022.10.13.512154

**Authors:** Meet Barot, Vladimir Gligorijevic, Richard Bonneau, Kyunghyun Cho

## Abstract

Knowledge of protein function is necessary for understanding biological systems, but the discovery of new sequences from high-throughput sequencing technologies far outpaces their functional characterization. Beyond the problem of assigning newly sequenced proteins to known functions, a more challenging issue is discovering novel protein functions. The space of possible functions becomes unlimited when considering designed proteins. Protein function prediction, as it is framed in the case of Gene Ontology term prediction, is a multilabel classification problem with a hierarchical label space. However, this framing does not provide guiding principles for discovering completely novel functions. Here we propose a neural machine translation model in order to generate descriptions of protein functions in natural language. In this way, instead of making predictions in a limited label space, our model generates descriptions in the language space, and thus is capable of composing novel functions. Given the novelty of our approach, we design metrics to evaluate the performance of our model: correctness, specificity and robustness. We provide results of our model in the zero-shot classification setting, scoring functional descriptions that the model has not seen before for proteins that have limited homology to those in the training set. Finally, we show generated function descriptions compared to ground truth descriptions for qualitative evaluation.

## 1 Introduction

Determining the function of proteins is a fundamental problem in biology. Accurately identifying these functions through wetlab experimentation is costly, so computational approaches to predict protein function have been necessary to reduce the functional search space for experimentalists. However, many existing approaches to protein function prediction are only able to predict known functional categories, leaving out the possibility of classifying proteins into new categories.

In this work, we propose a framing of the protein function prediction problem that does not rely on discrete categories. Instead, we directly predict the common functional description of a group of proteins in natural language, modeling the problem as a neural machine translation task. We train our model on about 300k protein sequences from the Swiss-Prot database [Bairoch and Apweiler, 2000] annotated with functional descriptions from the Gene Ontology (GO) [Ashburner et al., 2000]. We show that the model is capable of generating accurate function descriptions of proteins that are less than 30% identical to sequences in the training set and that have functions not present in the training set. We also propose three metrics to evaluate the correctness, specificity, and robustness of any model that can assign probabilities to a given sequence set and description.

## 2 Related Work

### 2.1 Protein Function Prediction

Many methods have been proposed for protein function prediction, though most do not consider the problem of discovering novel functions or generating their descriptions. As observed by Friedberg [2006], this has mainly been because of inherent difficulties of the flexibility of natural language, such as synonymous terms and ambiguity. These same difficulties were what led to the development of controlled and well-defined vocabularies of protein function, such as the Enzyme Commission Classification [Webb et al., 1992] and the Gene Ontology. As a result, the protein function prediction problem is generally framed as a supervised or semi-supervised multilabel classification problem with a structured output defined by these vocabularies, where the predicted labels are assumed to have some example in the training set [Bonetta and Valentino, 2020]. Much focus has been placed on this framing. The Critical Assessment of Functional Annotation (CAFA) [Zhou et al., 2019] serves as the main community benchmark for protein function prediction, and drives the field to improve upon previous methods. The CAFA evaluation considers proteins that can be described by existing categories. Yet many unlabeled proteins, especially in understudied organisms, are likely to perform functions that have not been seen before. The supervised approach does not address this possibility, and so new methods must be proposed for function discovery.

### 2.2 Clustering

Flat clustering-based approaches, by themselves, are not able to give much information about the new functional categories that they predict. They can only predict that a protein may belong to a category that has not been studied. One could compute average distances to clusters that contain known proteins, but beyond this, there is no testable hypothesis that the model can give about their function. NeXO [Dutkowski et al., 2013] and CliXO [Kramer et al., 2014] are both methods that generate an ontology of protein functions given relationships between proteins using hierarchical clustering. They aim at discovering novel functions. However, information about those new functions still rely on comparing the groupings to existing ontologies such as GO. Wang et al. [2018] describe a method that creates a concept hierarchy from phrases automatically extracted from scientific literature. This concept hierarchy is then aligned with the CliXO ontology in order to annotate proteins. However, this approach is still less flexible than generating free-form natural language.

### 2.3 Zero-shot learning approaches

Zero-shot learning approaches attempt to address the unseen class problem directly. DeepGOZero [Kulmanov and Hoehndorf, 2022] is a method that uses ontology axioms to predict for classes with no examples in the training set. However, the classes that are able to be predicted must be defined with ontological relations to seen classes. A similar limitation applies to clusDCA [Wang et al., 2015], which uses ontology relations to embed GO terms into a low dimensional space to perform zero-shot classification.

This constraint both restricts the possible novel functions that can be discovered and may not give sufficient information to design an experiment to test for the novel function.

### 2.4 Text generation and neural machine translation

Neural network-based text generation approaches have made significant progress in generating fluent and meaningful text [Fatima et al., 2022]. Further, deep learning-based techniques have shown promising results in image captioning methods [Hossain et al., 2019] and zero-shot classification of images[Radford et al., 2021]. Given enough data, deep learning methods have been shown to be capable of mapping between a range of input modalities and natural language. So far, there have been a few attempts to apply these methods to the protein function prediction domain. Zhang et al. [2020] use a graph-based generative model to generate Gene Ontology term names. However, the generation is limited to short phrases and relies on text descriptions from the GeneCards database [Safran et al., 2021] for the input.

Neural machine translation (NMT) is the automatic translation of written text from one natural language to another directly using neural networks [Cho et al., 2014]. NMT models have been widely deployed in production translation systems and show promise in domains other than natural language. Recently, a method called ProTranslator [Xu and Wang, 2022] has been proposed, which uses sequence, network and text description information concatenated into a 1-D feature vector in order to perform zero-shot classification on Gene Ontology terms. The authors also show that they are able to generate accurate and detailed descriptions for a set of proteins using a separate transformer model with this feature representation. Compared to ProTranslator, our method does not use any additional information to produce descriptions besides a set of protein sequences, and our model is trained directly to generate descriptions without pooling and losing positional information over the input sequences.

## 3 Methods

The following subsections give the motivation and formulations of the components of our method. Figure 1 contains a high-level overview.

**Figure 1:**
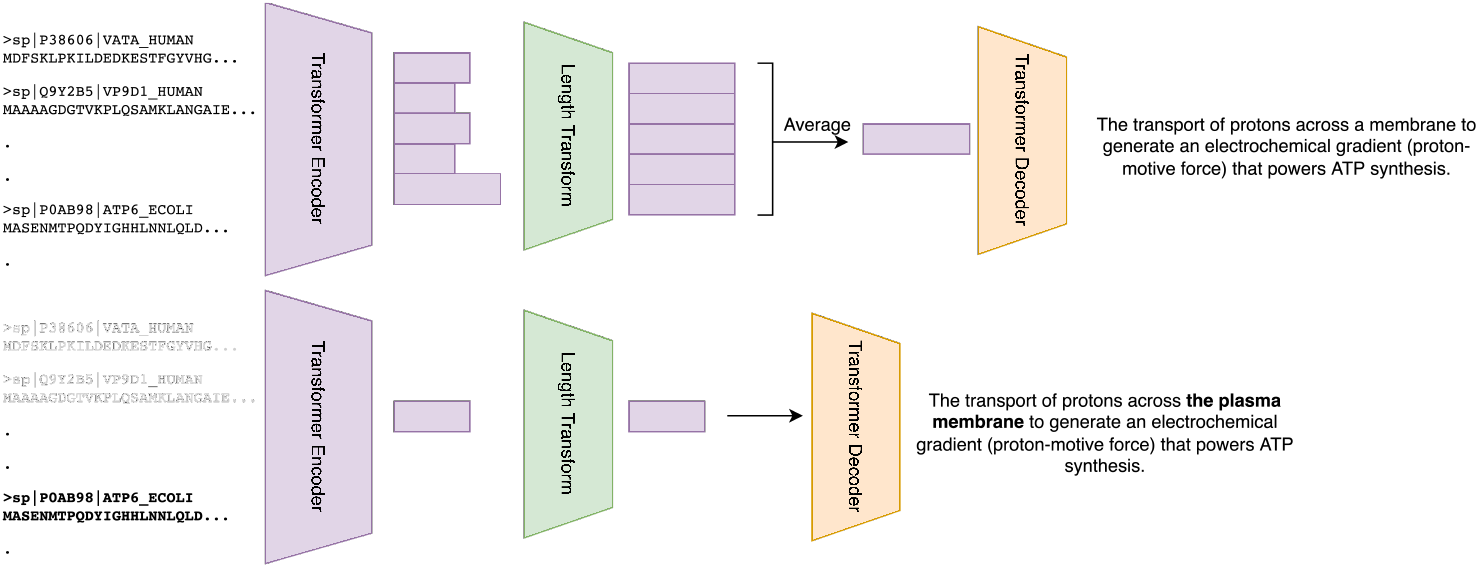
High-level diagram of the proposed transformer encoder-decoder model. The model is trained to produce the most specific common function of the input protein sequences.

### 3.1 Protein sets to describe

Biologists describe and categorize functions as abstractions of the common activity of a group of proteins, so we want our model to be able to perform this abstraction in a similar way. Formulating the problem as finding a single functional description for a single protein at a time is ill-defined, since a protein may have more than one distinct function Jeffery [2018]. Our task, then, is to find a description of the most specific function, *d_S_*, for a set of sequences, *S* = {*s*_1_, *s*_2_,…, *s_n_*}, of usually different lengths, |*s*_1_| = *l*_1_,…, |*s_n_*| = *l_n_*, that is common to all protein sequences *s_i_* ∈ *S*. There is still a possibility that there is more than one specific common function among the set, but it is less likely with larger sets, e.g., |*S*| =32.

### 3.2 Transformer encoder-decoder model with length transform

We use a transformer encoder-decoder model [Vaswani et al., 2017] with a length transform [Shu et al., 2020] to handle differing sequence lengths in order to average sequence features from the encoder. As a result of defining the learning task as a many-to-one problem, it was necessary to find a way to represent the common features of the set of sequences. The sequences’ representations should ideally be combined in some way that preserves amino acid ordering information, so we use the length transform in order to stretch the representations to the same shape in order to be averaged. This kind of length transform has been used previously in non-autoregressive neural machine translation problems Shu et al. [2020] and in protein design for changing protein sequence representations and generating sequences of variable lengths Gligorijevic et al. [2021]. For each sequence *s* ∈ *S*, we use a transformer model with positional encoding and self-attention to obtain a representation *h_s_* which consists of |*s*| continuous-valued vectors. As described in Shu et al. [2020], the length transform takes the input *h_s_* of length |*s*| and transforms the sequence with a monotonic location-based attention into 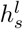 where *l* is the chosen output length so that 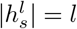. We choose *l* = *max*_*s*∈*S*_|*s*|.

### 3.3 Autoregressive generation of descriptions

It is desirable to represent protein function in a compositional way, so that the model has the ability to describe any given set of proteins without having to rely on examples of proteins with that specific function. To do this, we generate protein function descriptions in natural language, which gives the model the capability to compose a new function. We predict the tokens autoregressively, which is a standard practice in the NMT literature of top performing methods. With the |*S*| sequence representations 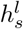 having all the same length after the length transform, we are able to take the average of these abstract representations, giving us *h_S_*, the representation of the whole sequence set. We use this representation in the transformer decoder in order to predict the next token of the description *d* given all the previous tokens.

### 3.4 Zero-shot Classification setting

Fundamentally, our model assigns probabilities to pairs of protein sets and descriptions. In order to evaluate the method, we use the zero-shot classification setting, where we wish to classify proteins into unseen categories. We develop three metrics in the Evaluation section to evaluate the conditional probability distribution *P*(*d_S_*|*S*) learned by the model in this classification setting.

### 3.5 Generation (beam search)

Generation of descriptions is a search problem through the set of all possible output token sequences, where the goal is to find the sequence with the largest probability. Generation given an autoregressive model is a highly studied problem in the natural language processing literature. We use beam search Graves [2012] in the current implementation in order to find reasonable generated descriptions. We use a beam width of 10 with a length penalty of 2.0. Direct evaluation of these descriptions is an unsolved problem: currently, manual inspection by expert human evaluators is the best method we have.

## 4 Evaluation

In this section, we define three metrics that can be computed using known functional descriptions in order to evaluate our models’ learned probability distributions.

Generated descriptions are shown in the Results section for qualitative analysis. Quantitative analysis of the generated descriptions requires data from human evaluators with expertise in protein function in order to determine the accuracy of generated descriptions. A framework for performing that analysis with expert curators is explored in the Discussion section.

### 4.1 Attribute 1: Annotation correctness

Given a sequence set for which the model is assigning scores to function descriptions, descriptions of GO terms that annotate the entire sequence set should be scored higher than terms that do not annotate the entire sequence set.

Let *D_S_* be the GO term descriptions associated with sequence set S.

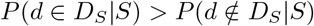

A way to measure this attribute would be to calculate:

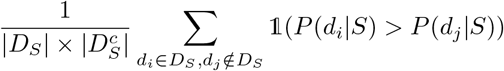

where 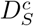 is the complement of *D_S_* and 1 is the indicator function.

### 4.2 Attribute 2: Specificity preference

Among terms that do annotate the whole set, the model should score child terms higher than their ancestor terms. Let *A*(*d*) denote the description of a direct parent of the GO term described by *d.*

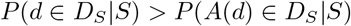

Note: any protein set that is annotated with d would always be annotated with *A*(*d*), *A*(*A*((*d*)) and so on.

A way to measure this attribute would be to calculate:

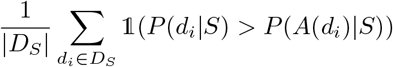

### 4.3 Attribute 3: Annotation robustness

Any set of sequences that have the same exact set of GO descriptions in common should be scored with the same rankings for those GO descriptions.

Let *S_i_* and *S_j_* be different sequence sets such that *D_S_i__* = *D_S_j__* and *S_i_* ≠ *S_j_*, and let *R*(*X*) be a ranking function that gives the ranks of entries in *X*, in descending order.

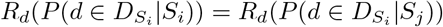

A way to measure this attribute would be to calculate the average Spearman’s rank correlation of the rankings for all sequence sets’ correct descriptions. Let *R_S_i__* = *R*(*P*(*D_S_i__*|*S_i_*)):

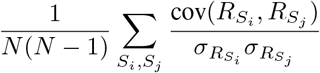

where *N* is the total number of sequence sets that have the exact set of GO descriptions *D_S_i__*. In reality, this number may be too large to actually sum (especially if |*D_S_i__*| is small), so we approximate this measure by subsampling *n* < *N* sequence sets to average over instead. The sum is only calculated over non-identical pairs of sequence sets.

## 5 Data

We take sequences and annotations from the Uniprot-KB Swiss-Prot database, which is manually annotated and reviewed, in order to create our training and evaluation sets of proteins and function descriptions. This database had 566,996 proteins total. To show that our model can generalize to non-homologous proteins, we clustered the proteins into groupings with less than 30% sequence identity using cd-hit [Li and Godzik, 2006], and separated these into training and test sets. To focus on the functions that were both specific enough and had a sufficient number of examples in our evaluation sets, we restricted the maximum number of proteins per GO term to 1280, and minimum number of proteins to 32. Hyperparameters chosen were tuned on the training set proteins with training function descriptions. The number of proteins and GO terms that were used after these restrictions in our training set and evaluation sets are listed in Table 1.

**Table 1:** Number of proteins and GO terms in training and test sets.

|  | Train P&F | Train P, Test F | Test P, Train F | Test P&F |
| --- | --- | --- | --- | --- |
| Proteins | 316k | 181k | 20k | 20k |
| GO terms | 9k | 2k | 879 | 1.5k |

## 6 Results

We show model performances in Table 2. The table suggests that the model is able to rank unseen functions for protein sets that it has been exposed to in training, with the model’s rankings of identically annotated sets being in moderate agreement. For test proteins that have less than 30% sequence identity to the training set, the model is still able to assign rankings of 1000 randomly selected functions from the training set with a correctness 30% above random assignment (0.5). For the low-similarity test proteins that have functions that are not seen in the training set, the model is still able to rank 21% better than random rankings.

**Table 2:**
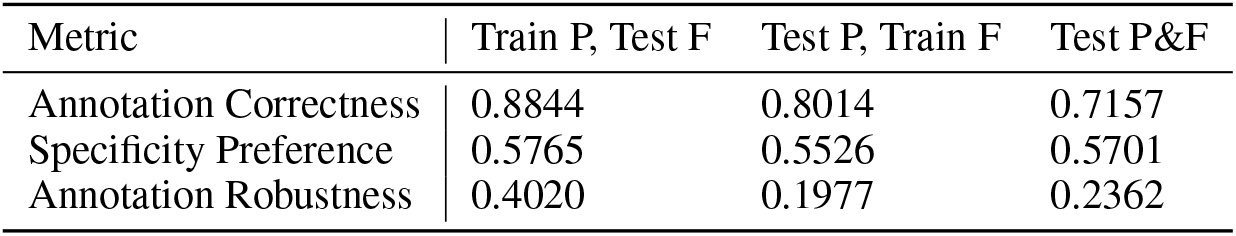
Model Performances

We are mainly focused on using the model for generation, and these metrics are meant mostly as guides for model design. The loss function used is not optimizing for classification accuracy; it is optimizing the model’s probability distribution to assign high probability to descriptions assigned to a sequence set.

We show sample test set descriptions in Table 3. The first row shows that the model describes verbatim a related term (GO:0001654, eye development) for the proteins selected, whereas the true term is appendage development (GO:0048736). Their common ancestor term is anatomical structure development (GO:0048856). This description is more specific than the actual term from which the proteins are sampled, but it is not accurate. The next generated description is more general than the actual description of the sampled set (modulates vs. activates), but is correct; it is the direct parent of the true term. The third generated description is related but ultimately different than the actual description of the protein set. The fourth generated description is more specific than both the true common GO description of the set (protein import, GO:0017038) and the generated description’s closest known GO term, protein exit from endoplasmic reticulum (GO:0032527). It is describing protein import into the nucleus from the endoplasmic reticulum, which is not currently a GO term, but if it was, it would be a descendant of both of these terms.

**Table 3:** Sample Test Set Description Generations

| True Common GO Term | True Common GO Description of Sequence Set | Model Generated Description of Sequence Set | Closest Known GO Term to Generated Description |
| --- | --- | --- | --- |
| GO:0048736, appendage development | <SOS> the process in which the anatomical structures of appendages are generated and organized . an appendage is an organ or part that is attached to the trunk of an organism . <EOS> | <SOS> the process whose specific outcome is the progression of the eye over time , from its formation to the mature structure . <EOS> | GO:0001654, eye development |
| GO:0045597, positive regulation of cell differentiation | <SOS> any process that activates or increases the frequency , rate or extent of cell differentiation . <EOS> | <SOS> any process that modulates the frequency , rate or extent of cell differentiation . <EOS> | GO:0045595, regulation of cell differentiation |
| GO:0071162, CMG complex | <SOS> a protein complex that contains the gins complex , cdc45p , and the heterohexameric mcm complex , and that is involved in unwinding dna during replication . <EOS> | <SOS> any process involved in forming the mature 3 ' end of a dna ( mrna ) molecule . <EOS> | GO:0031124, mRNA 3'-end processing |
| GO:0017038, protein import | <SOS> the targeting and directed movement of proteins into a cell or organelle . not all import involves an initial targeting event . <EOS> | <SOS> the directed movement of proteins from endoplasmic reticulum to the nucleus . <EOS> | GO:0032527, protein exit from endoplasmic reticulum |

## 7 Discussion

In this work, we have proposed a novel method to generate protein function descriptions in order to discover new protein functions. We have demonstrated that our model can accurately rank unseen function descriptions for proteins not seen in the training set, and show promising results in generated function descriptions. Given that this model is trained using raw text descriptions of protein function, it is possible to extend this work to use descriptions from other databases besides the Gene Ontology, such as Pfam [Bateman et al., 2004], KEGG[Kanehisa et al., 2002], or Enzyme Commission classes. This increase in data could allow for higher quality descriptions, or the ability to query the model to output descriptions of a particular aspect of function. Below, we explore how we might further evaluate the method’s generated descriptions using human expertise and curation.

### 7.1 Future human-assisted evaluation of function discovery

As our scoring metrics for evaluation are automated, they can be used for optimizing the architecture and other hyperparameters of the model (either manually or with some search method). However, in the case of actual use on proteins that are not very well studied, it can be difficult to know whether a given description is accurate. Human-assisted evaluation will be needed for the descriptions generated for a given set of novel proteins. This feedback could be used to fine-tune the model to produce more accurate, fluid or generally desirable descriptions of proteins, as has been done for document summarization models [Ziegler et al., 2019, Stiennon et al., 2020].

One possible way of obtaining human feedback would be to ask an expert with knowledge of the Gene Ontology and familiarity with some families of proteins to choose between two descriptions for a given sequence set that is generated from a trained model. Doing this over a large enough dataset would allow us to train a reward estimation model that can then be used to fine-tune the original trained model using reinforcement learning. However, this would be expensive, as the task needs to be done by an expert. Richer information, such as ranking the similarities to an existing GO term, or suggesting changes to particular portions of the description, could be used to increase performance even with a small number of examples with human feedback.

## References

Amos Bairoch and Rolf Apweiler. The swiss-prot protein sequence database and its supplement trembl in 2000. Nucleic acids research, 28(1):45–48, 2000.

Michael Ashburner, Catherine A Ball, Judith A Blake, David Botstein, Heather Butler, J Michael Cherry, Allan P Davis, Kara Dolinski, Selina S Dwight, Janan T Eppig, et al. Gene ontology: tool for the unification of biology. Nature genetics, 25(1):25–29, 2000.

Iddo Friedberg. Automated protein function prediction—the genomic challenge. Briefings in bioinformatics, 7(3):225–242, 2006.

Edwin C Webb et al. Enzyme nomenclature 1992. Recommendations of the Nomenclature Committee of the International Union of Biochemistry and Molecular Biology on the Nomenclature and Classification of Enzymes. Number Ed. 6. Academic Press, 1992.

Rosalin Bonetta and Gianluca Valentino. Machine learning techniques for protein function prediction. Proteins: Structure, Function, and Bioinformatics, 88(3):397–413, 2020.

Naihui Zhou, Yuxiang Jiang, Timothy R Bergquist, Alexandra J Lee, Balint Z Kacsoh, Alex W Crocker, Kimberley A Lewis, George Georghiou, Huy N Nguyen, Md Nafiz Hamid, et al. The cafa challenge reports improved protein function prediction and new functional annotations for hundreds of genes through experimental screens. Genome biology, 20(1):1–23, 2019.

Janusz Dutkowski, Michael Kramer, Michal A Surma, Rama Balakrishnan, J Michael Cherry, Nevan J Krogan, and Trey Ideker. A gene ontology inferred from molecular networks. Nature biotechnology, 31(1):38–45, 2013.

Michael Kramer, Janusz Dutkowski, Michael Yu, Vineet Bafna, and Trey Ideker. Inferring gene ontologies from pairwise similarity data. Bioinformatics, 30(12):i34–i42, 2014.

Sheng Wang, Jianzhu Ma, Michael Ku Yu, Fan Zheng, Edward W Huang, Jiawei Han, Jian Peng, and Trey Ideker. Annotating gene sets by mining large literature collections with protein networks. In Pacific Symposium On Biocomputing 2018: Proceedings of the Pacific Symposium, pages 602–613. World Scientific, 2018.

Maxat Kulmanov and Robert Hoehndorf. Deepgozero: Improving protein function prediction from sequence and zero-shot learning based on ontology axioms. bioRxiv, 2022. doi: 10.1101/2022.01.14.476325. URLhttps://www.biorxiv.org/content/early/2022/01/14/2022.01.14.476325.

Sheng Wang, Hyunghoon Cho, ChengXiang Zhai, Bonnie Berger, and Jian Peng. Exploiting ontology graph for predicting sparsely annotated gene function. Bioinformatics, 31(12):i357–i364, 2015.

Noureen Fatima, Ali Shariq Imran, Zenun Kastrati, Sher Muhammad Daudpota, Abdullah Soomro, and Sarang Shaikh. A systematic literature review on text generation using deep neural network models. IEEE Access, 2022.

MD Zakir Hossain, Ferdous Sohel, Mohd Fairuz Shiratuddin, and Hamid Laga. A comprehensive survey of deep learning for image captioning. ACM Computing Surveys (CsUR), 51(6):1–36, 2019.

Alec Radford, Jong Wook Kim, Chris Hallacy, Aditya Ramesh, Gabriel Goh, Sandhini Agarwal, Girish Sastry, Amanda Askell, Pamela Mishkin, Jack Clark, et al. Learning transferable visual models from natural language supervision. In International Conference on Machine Learning, pages 8748–8763. PMLR, 2021.

Yanjian Zhang, Qin Chen, Yiteng Zhang, Zhongyu Wei, Yixu Gao, Jiajie Peng, Zengfeng Huang, Weijian Sun, and Xuan-Jing Huang. Automatic term name generation for gene ontology: task and dataset. In Findings of the Association for Computational Linguistics: EMNLP 2020, pages 4705–4710, 2020.

Marilyn Safran, Naomi Rosen, Michal Twik, Ruth BarShir, Tsippi Iny Stein, Dvir Dahary, Simon Fishilevich, and Doron Lancet. The genecards suite. In Practical guide to life science databases, pages 27–56. Springer, 2021.

Kyunghyun Cho, Bart Van Merriënboer, Dzmitry Bahdanau, and Yoshua Bengio. On the properties of neural machine translation: Encoder-decoder approaches. arXiv preprint arXiv:1409.1259, 2014.

Hanwen Xu and Sheng Wang. Protranslator: zero-shot protein function prediction using textual description. In International Conference on Research in Computational Molecular Biology, pages 279–294. Springer, 2022.

Constance J Jeffery. Protein moonlighting: what is it, and why is it important? Philosophical Transactions of the Royal Society B: Biological Sciences, 373(1738):20160523, 2018.

Ashish Vaswani, Noam Shazeer, Niki Parmar, Jakob Uszkoreit, Llion Jones, Aidan N Gomez, Łukasz Kaiser, and Illia Polosukhin. Attention is all you need. Advances in neural information processing systems, 30, 2017.

Raphael Shu, Jason Lee, Hideki Nakayama, and Kyunghyun Cho. Latent-variable non-autoregressive neural machine translation with deterministic inference using a delta posterior. In Proceedings of the AAAI Conference on Artificial Intelligence, volume 34, pages 8846–8853, 2020.

Vladimir Gligorijevic, Daniel Berenberg, Stephen Ra, Andrew Watkins, Simon Kelow, Kyunghyun Cho, and Richard Bonneau. Function-guided protein design by deep manifold sampling. bioRxiv, 2021.

Alex Graves. Sequence transduction with recurrent neural networks. arXiv preprint arXiv:1211.3711, 2012.

Weizhong Li and Adam Godzik. Cd-hit: a fast program for clustering and comparing large sets of protein or nucleotide sequences. Bioinformatics, 22(13):1658–1659, 05 2006. ISSN 1367-4803. doi: 10.1093/bioinformatics/btl158.

Alex Bateman, Lachlan Coin, Richard Durbin, Robert D Finn, Volker Hollich, Sam Griffiths-Jones, Ajay Khanna, Mhairi Marshall, Simon Moxon, Erik LL Sonnhammer, et al. The pfam protein families database. Nucleic acids research, 32(suppl_1):D138–D141, 2004.

Minoru Kanehisa et al. The kegg database. In Novartis foundation symposium, pages 91–100. Wiley Online Library, 2002.

Daniel M Ziegler, Nisan Stiennon, Jeffrey Wu, Tom B Brown, Alec Radford, Dario Amodei, Paul Christiano, and Geoffrey Irving. Fine-tuning language models from human preferences. arXiv preprint arXiv:1909.08593, 2019.

Nisan Stiennon, Long Ouyang, Jeffrey Wu, Daniel Ziegler, Ryan Lowe, Chelsea Voss, Alec Radford, Dario Amodei, and Paul F Christiano. Learning to summarize with human feedback. Advances in Neural Information Processing Systems, 33:3008–3021, 2020.

